# Postural control in an upright snake

**DOI:** 10.1101/2024.12.13.628369

**Authors:** Ludwig A. Hoffmann, Petur Bryde, Ian C. Davenport, S Ganga Prasath, Bruce C. Jayne, L Mahadevan

## Abstract

Posture and its control are fundamental aspects of animal behavior that capture the complex interplay between sensorimotor activity driven by muscular forces and mediated by environmental feedback. An extreme example of this is seen in brown tree snakes and juvenile pythons, which can stand almost upright, with 70% of their body length in the air. We quantify experimental observations of this behavior and present a minimal theoretical framework for postural stability by modeling the snake as an active elastic filament whose shape is controlled by muscular forces. We explore two approaches to characterize the musculature needed to achieve a specific posture: proprioceptive feedback (whereby the snake senses and reacts to its own shape) and a control-theoretic optimization approach (whereby the snake minimizes the expended energy to stand up), and also analyze the dynamic stability of the snake in its upright pose. Our results lead to a three-dimensional postural stability diagram in terms of muscle extent and strength, and gravity, consistent with experimental observations. In addition to general predictions about posture control in animals, our study suggests design principles for robotic mimics.

## I. INTRODUCTION

If snakes on a plane [1] are scary, just imagine snakes out-of-the-plane, navigating from one tree branch to another, like Kaa in *The Jungle Book* [2], posturing to strike fear and awe. Pose and posture are crucial aspects of behavior in animals [3, 4]; they allow for function, e.g. seeing further, or displays that serve as signals and as a precursor to dynamic maneuvers. Controlling posture requires a sense of one’s own shape and orientation (proprioception) that allows an animal to dynamically react to environmental stimuli and fluctuations. This results in a range of different strategies, from swimming fish that sense their body shape [5, 6] to human bipedalism being (partially) controlled by a sense of equilibrium and one’s own posture [7–9].

Snakes and their postures stand out among vertebrates as rather unique because of their uniform slender geometry, an absence of limbs, and an extreme ability to deform their body through muscular actuation. Indeed, since snakes are found in a wide variety of terrestrial, arboreal, aquatic, and even aerial habitats, their poses, postures and modes of locomotion reflect the challenges associated with traversing these diverse environments. Compared to a wealth of recent studies on the modes of snake locomotion and their implications for creating artificial mimics [10–12], their poses and postures are poorly understood. Recent experiments [3, 13] have highlighted the role of muscle activity and postures of snakes bridging various gaps. Snakes perform this reaching task by selectively activating the epaxial muscles along vertebrae to control stiffness and prevent mechanical instabilities such as self-buckling. When the pitch and yaw angles required to traverse a gap vary, snakes use different axial motor patterns [13], and snakes can bridge much greater gap distances going straight up rather than horizontally [14]. Two theoretical possibilities for supporting the suspended portion of the body are simultaneously activating muscular antagonists to make the body rigid (presumably uneconomical) or only activating the muscles required to create a particular shape and oppose the effects of gravity that would disrupt that shape (economical and requiring minimal effort). Inspired by these observations and different possibilities for motor control, we develop a model to describe and predict the shape and muscular control that would require minimal effort for extreme posture in snakes standing almost upright with most of their body in the air, see Fig. 1.

**Figure 1.**
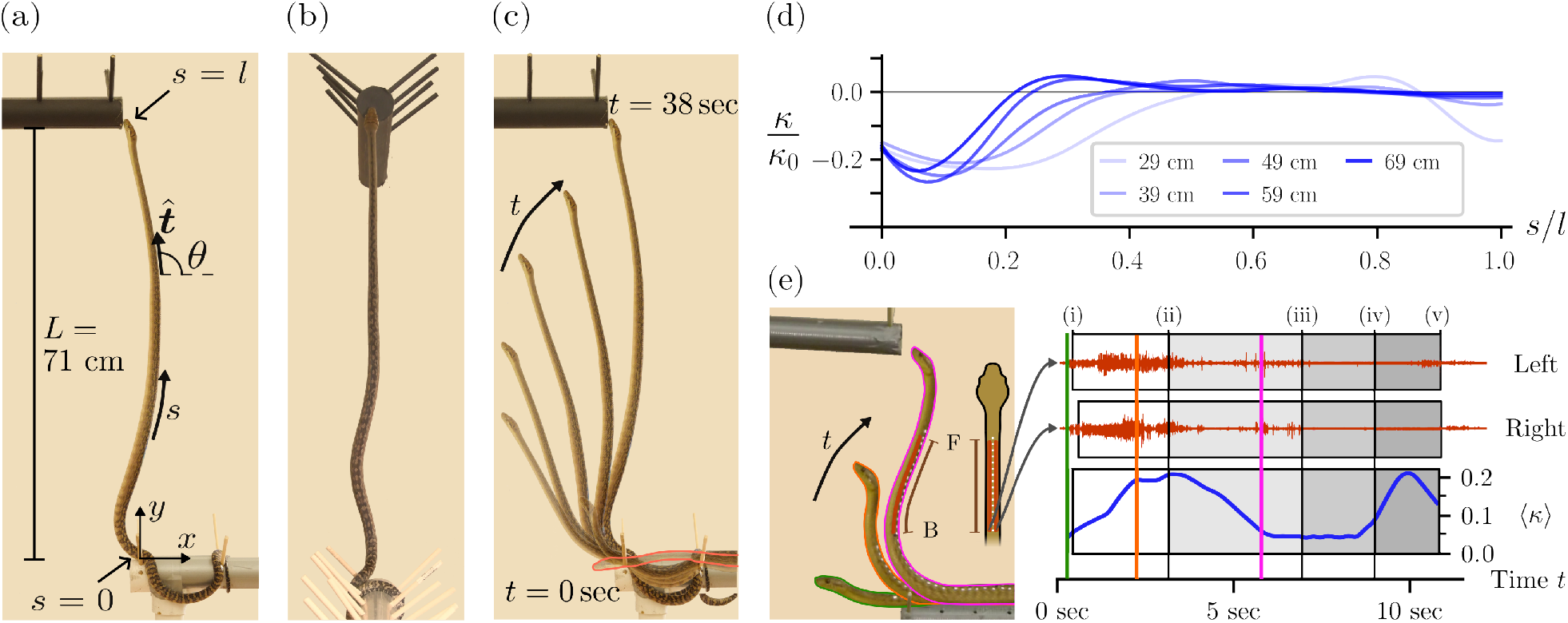
Posture of standing snake. (a) Lateral and (b) dorsal view of a scrub python which is lifting itself up while “standing” on a perch. The position is shortly before the snake contacts the upper perch at height *L* = 0.71 m, which corresponds to 70% of the snakes’ total length. *s* is the coordinate along the center-line of the part of the snake’s body that is off the perch, with *s* = 0 the point of contact with the lower perch and *s* = *l* the head. (c) Overlay of frames of the snake taken at different times. Earlier times are shown more transparent. At *t* = 0 sec (outlined with red) the snake’s head first leaves the perch while the last image (*t* = 38 sec) is just before the snake touches the upper perch. The time between successive images is approximately 6.3 seconds. (d) Lateral-view curvature *κ(s)* of the python’s body at different times where the length of the body in the air increases from *l* = 0.29 m to *l* = 0.69 m. The curvature is normalized by dividing by *κ*_0_ = 1/1.9 cm, the inverse diameter of the snake. Negative values of curvature indicate regions that are concave dorsally. The curvatures shown are an average over four independent trials with the same snake. As the total time of each trial varies we use *l*, rather than time, as a measure of progress. (e) Activity of the semispinalis-spinalis (SSP) muscles for a brown tree snake standing up. The SSP muscles are located dorsally on the left and right side of the neural spines of the vertebrae (indicated in red in the top-view sketch and one of the snake images). They function as dorsiflexors and arch the back when contracting bilaterally while inducing lateral flexion when activated unilaterally. The activity of the muscle segments in the location between points F and B (indicated by the brown line) is measured using two electrodes inserted into the left and right SSP muscles near point B (see sketch). The activity over time is shown in the electromyograms on the right (top two graphs). The outline colors of the three shapes on the left correspond to the respective times marked by the three colored lines on the right. At points (i) and (ii), F and B leave the lower perch, while at (iii) the snake’s head reaches the upper perch. Points F and B reach the upper perch at (iv) and (v), respectively. We compute ⟨ *κ*(t) ⟩, defined as the integral of |*κ*(*s, t*)| over the length between F and B and normalized by the snake’s diameter, to measure average curvature (bottom graph on the right). Initially, when the region of interest first bends, muscular activity is high and symmetrical between the left and right sides. Once the curved region is established, muscular activity decreases significantly. When the snake reaches the upper perch, curvature increases without a corresponding rise in muscular activity. Electromyogram data adapted from Ref. [13].

## II. EXPERIMENTAL OBSERVATIONS

For two species that readily climb trees (brown tree snake, *Boiga irregularis* [*N* = 3]; scrub python, *Simalia amesthistina* [*N* = 1]), we elicited gap bridging between two horizontal pipes which were a vertical distance *L* apart. Each 5 cm diameter perch had several pairs of pegs (length = 10 cm; diameter = 0.5 cm) placed at 10 cm intervals (Fig. 1). Each trial started by placing a snake on the lower perch, and we analyze the movements from when the head passed the end of the lower perch until the snake reached the upper perch. The distance *L* was increased progressively from 30 cm to 80 cm or until the snakes were unable to bridge the gap (see SI Sec. S3 for details). Remarkably, the brown tree snakes and scrub python could reach upper perches even when the distance *L* exceeded 50% and 70%, respectively, of their total body length *l*_tot_ (see Fig. 1(a, b) for snapshots of lateral and dorsal view, SI Figs. S2, S3, and SI Movies 1-4). A time-series of the climbing process is shown in Fig. 1(c). For large *L*, small oscillations occurred towards the end of the process (see SI Movies 1-4). However, these do not significantly affect the shape of the snake observed in lateral view, and we will postpone discussing them below to Sec. III C. First, we are interested in investigating how a snake’s shape changes during the standing-up, and in isolating general features of this process. While the lateral-view shape is found to be universal across all trials, the dorsal-view shape is not, but is instead related to the position of the snake on the lower perch (shown in SI Fig. S3). Therefore, even though the dorsal-view curvature does not vanish, we focus on analyzing only the lateral-view shape. To quantify the standing-up process we extract the shape of the snake’s body for each successive frame (30 frames/sec) of the video of a chosen trial and compute the lateral curvature *κ*(*s, t*), where *s* is the arc-length along the snake’s body and *t* the elapsed time from when the head of the snake first left the lower perch (ref. Fig. 1(a) and Sec. S2 for details, see also SI Movies 3,4 and Tab. S1 for a list of variables). Fig. 1(d) shows the resulting profile *κ*(*s, t*)*/k*_0_ vs *s/l* at different times *t* for the scrub python with *L* = 0.7*l*_tot_. Here, *l* is the length of the snake’s body in the air and *κ*_0_ is chosen to be the inverse diameter of the snake; see also SI Fig. S2 for an analogous representative example of a brown tree snake. Note that the curvature shown in Fig. 1(d) is the average value over 4 trials for the same snake and at fixed *L* = 0.71 m indicating that the strategy for climbing is reproducible (particularly at later times, see SI Fig. S4).

As the total duration of the process varies across trials we use the length *l*, rather than time, as a measure of the climbing progress (described in detail in SI Sec. S3). We observe that during the climbing process the snake’s shape is characterized by a boundary layer near the lower perch (*s* = 0) where curvature is concentrated, and outside of which the curvature is relatively small, typically vanishing near the snout (at *s* = *l*). Once established, the boundary layer location is approximately constant in time (as shown in Fig. 1(d)). Furthermore, while the variations in shape over trials are large for early times, at late times we observe a convergence towards a universal shape, with minimal variance (SI Fig. S4).

These observations raise the following questions about posture and stability, respectively: (*i*) How to explain the “S-shape” of the standing snake in the sagittal plane as well as the existence of a boundary layer away from which the curvature vanishes?; (*ii*) How does the snake stabilize itself and prevent falling over despite up to 70% of its total body length being lifted up? In the following Sec. III, we develop a minimal theoretical model that we will use to describe the shape of a snake and investigate whether our model can describe the characteristic features of the snake’s posture and its stability.

## III. THEORY

As mentioned earlier, while we observe non-vanishing curvature in both sagittal as well as coronal plane, we focus on the sagittal plane as the shape dynamics in this plane are consistent across all experiments. Since the snake changes its posture relatively slowly (inertial effects are small compared to elastic effects), we consider the postural changes as occurring quasi-statically. This allows us to use the parametrization of the snake in terms of its center line ***r***(*s*) = (*x*(*s*), *y*(*s*)), where *s* is the arc length along the center-line of the snake’s body such that *s* = 0 is the point of contact with the lower perch and *s* = *l* is the position of the snake’s snout which is a slowly changing variable (shown in Fig. 1(a)). The unit tangent to the curve at point *s* is defined as,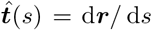 and can be written as 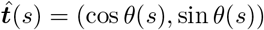 with θ(*s*) the angle between,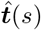 and the horizontal. The snake experiences an inhomogeneous force ***F*** (*s*) = (*F*_*x*_(*s*), *F*_*y*_(*s*)) and a moment *m*(*s*) along its body axis. Assuming an additive decomposition of the moment as the sum a passive moment *m*_p_(*s*) and an active component, *m*_a_(*s*) we write *m*(*s*) = *m*_p_(*s*)+ *m*_a_(*s*). The passive elasticity that resists curvature can be accounted for via the relation *m*_p_(*s*) = *B*_p_*κ(s)*, where *B*_p_ is the bending stiffness (assumed to be independent of *s*, as the snake’s geometry is approximately uniform) and *κ(s)* = d*θ*(*s*)*/* d*s* is the local curvature. In Fig. 1(e) we reproduce electromyography measurements from Ref. [13] which show that the actuation of the snake’s semispinalis-spinalis (SSP) muscles plays a vital role while bridging gaps, and in particular that the SSP muscles are activated mainly when the body is maximally bent, a fact that we will use later. Given the active and passive contributions to the moment at every cross-section, we can write the force and torque balance equations for the snake as [15]:

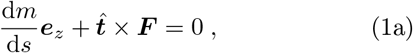

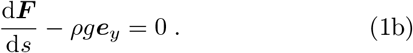

Here, ρ is the linear mass density and *g* is the gravitational constant. In order to solve for the shape of the snake described by the center-line, the force and moment balance equations need to be complemented by boundary conditions. Since the head of the snake is in the air, it must be free of forces, so that we have ***F*** (*s* = *l*) = 0. Using this condition while integrating Eq. (1b) yields *F*_*x*_(*s*) = 0 and *F*_*y*_(*s*) = − ρ*g*(*l* − *s*). Combining this with Eq. (1a), we find that:

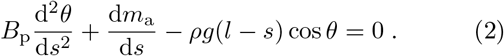

In the absence of active muscular forces and torques, *m*_a_ = 0 and the equation reduces to the classical equation of an elastic filament in a gravitational field [15], with a well known instability to self-buckling when the filament is tall/heavy enough [16]. An active snake can stabilize itself by varying *m*_a_, and from now on we will consider the single Equation (2) to investigate the posture of a standing snake, completely characterized by the angle *θ*(*s*), since (*dx/ds, dy/ds*) = (cos *θ*, sin *θ*). Here, we have limited ourselves to movements in the sagittal plane, consistent with observations.

### A. Proportional muscular activation of posture

Snakes (and other animals) are known to process information about their own shape via proprioception, the sense of their own body. They use this to control muscular activation and achieve a desired posture. In our minimal setting, this implies that *m*_a_ is a function of *κ*(*s, t*), since the shape of a planar filament is known to be completely characterized by its curvature (up to rigid motions). However, since muscles can have a delayed response to stimuli due to neuromechanical processes and have long-range connectivity with fibers that extend along many vertebrae, the curvature at a given position and time (*s, t*) can influence the muscular response at a different position and time (*s*^*′*^, *t*^*′*^), so that the most general linear relation connecting the two is

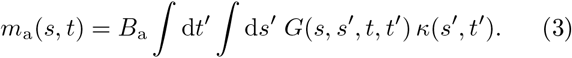

Here, *B*_a_ is a constant representing the *active bending stiffness* and *G*(*s, t*) is the kernel capturing potential non-locality and time-delay effects [5, 17]. We note that this relation is a natural generalization of linear response theory linking stimulus and response, and used widely to model many different systems, from bacteria to engineered control systems, applied here to the curvature.

For simplicity, we begin by analyzing the limit when the muscular activation kernel is local in time and space, *G*(*s, t*) = δ(*s*)δ(*t*). Eq. (3) simplifies and the active moment is directly proportional to the curvature, *m*_a_(*s*) = *B*_a_*κ(s)*, just as for the passive moment. Equation (2) then reads, after non-dimensionalization,

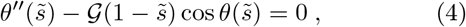

with the single non-dimensional parameter 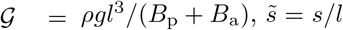 the arc-length parametrization, and primes denote differentiation with respect to 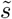. Unsurprisingly, the introduction of a local active moment is mathematically equivalent to augmenting the passive bending stiffness *B*_p_ → *B*_p_ +*B*_a_. In order to solve Eq. (4) we need to specify the boundary conditions for θ(*s*).

Our experiments show that at the perched end of the snake, which we choose to be the origin with *x*(0) = *y*(0) = 0, the snake is horizontal, thus *θ*(0) = π. The free snout of the snake at *s* = *l* shows different behaviors as more and more of the snake is in the air. Initially the snake moves horizontally off the perch, but only after a sufficient fraction is not in contact with the perch anymore, does it start to face upwards, presumably because the “free-plank” pose is hard to sustain as more and more of the snake is in the air [13]. We focus only on the latter and divide this behavior into two phases, a transient lift-up pose and a static standing-up pose. During lift-up (not shown in Fig. 2) the angle of the snout, *θ*(*l*), decreases from its initial value *θ*_init_ = π to a value *θ*_fin_ π*/*2, i.e., the body bends upwards while the extent of the snake in the air *l* does not increase significantly. On the other hand, in the standing-up phase (Fig. 2), *θ*(*l*) ≈ *θ*_fin_ remains approximately constant, while the length *l* changes significantly. Experimentally, we find that the dynamics of the initial lift-up phase vary significantly between different trials (see SI Movies 1-4) whereas the standing-up phase is quantitatively similar for different runs for the same snake and also qualitatively similar for different snakes. We therefore focus on theoretically describing the standing-up behavior (see the SI Sec. S4 for a more detailed discussion and theoretical results on the lift-up behavior). We set *θ*_fin_ = π*/*2 + 0.2 in the following, motivated by the experimental observation that the angle is slightly larger than the vertical. However, it is important to note that the selection of the exact value is somewhat arbitrary, as it can vary slightly both during individual experiments and between different trials.

**Figure 2.**
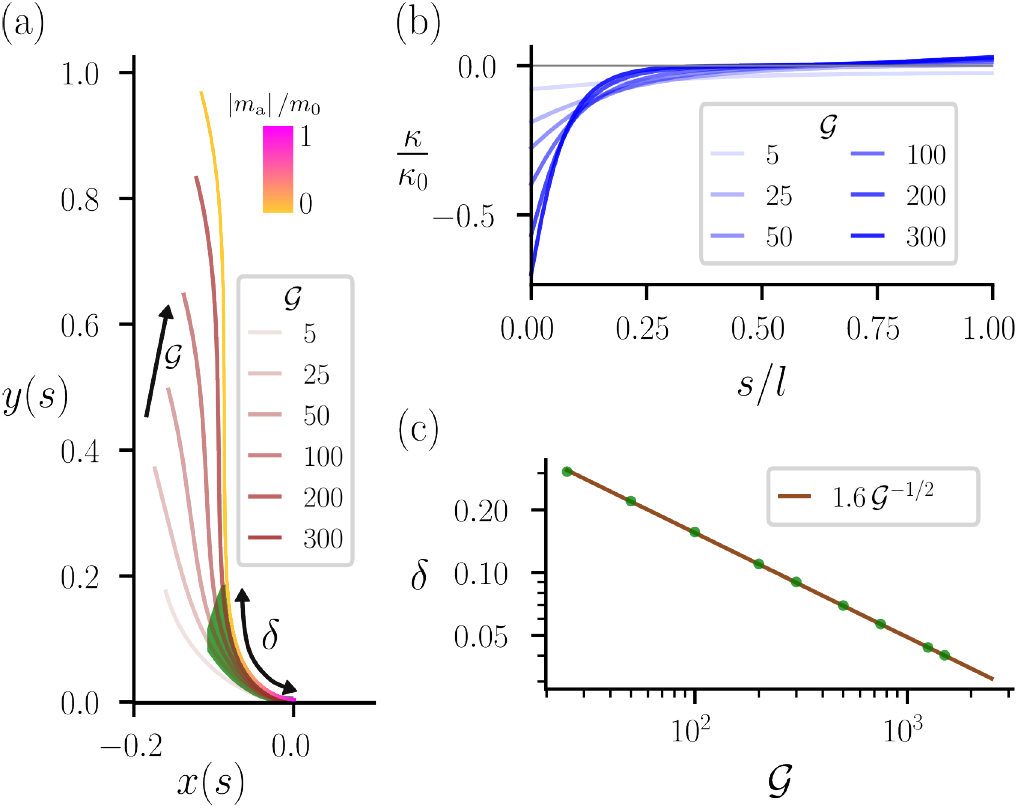
Posture for local, proportional feedback. (a) Shape found from numerically solving Eq. (4) for different values of the dimensionless parameter 𝒢 = *ρgl*^3^*/*(*B*_p_ + *B*_a_). 𝒢 increases from left to right, modeling the experimentally observed standing-up behavior where *l* increases while *θ*(*s* = *l*) is approximately constant. Here, we assumed that *B*_p_ + *B*_a_ remains constant during the process in order to rescale length scales by *l*, with *l* = 1 for 𝒢 = 300. *δ* denotes the boundary region defined as *κ*(*s/l* = *δ*) = *κ*(*s* = 0)*/*4, which is indicated by a green shading. We used the same normalization value *κ*_0_ as in Fig. 1. Moreover, we visualize the normalized muscular actuation |*m*_a_(*s*)| */m*_0_ along the curve for 𝒢 = 300 using a color map. (b) Curvature *κ(s)* of the shapes shown in (a) for different values of 𝒢. Note the existence of a boundary layer for small values of *s* and larger values of 𝒢, outside of which the curvature is small. (c) The length of the boundary layer *δ* for different values of 𝒢 (green points) and the best fit (brown line) of the theoretically predicted 𝒢^—1*/*2^ scaling (see main text). Note that while the prefactor of the fit changes for different definitions of *δ*, the scaling does not.

Numerical solutions of Eq. (4) during the standing-up phase (with boundary condition *θ*(*l*) = *θ*_fin_ fixed) for increasing values of the scaled gravity parameter 𝒢 ∼ *l*^3^, are shown in Fig. 2, and capture the climbing process as the unsupported length *l* increases. For large 𝒢 the shape of the curve converges towards the S-shape observed in experiments (see Fig. 1(c), SI Fig. S2), and reproduces the observation that, as the length of the snake in the air *l* increases, the change of shape of the snake is minimal. We also note that as 𝒢 increases, the curvature (and activity) localizes at a boundary layer near the perch *s* = 0 (see Fig. 2(b)); outside this boundary layer the curvature is very small, similar to the experimental measurements shown in Fig. 1(d), SI Fig. S2. A simple scaling analysis allows us to determine the extent of the boundary layer. First, we note that for 𝒢 ≫ 1 in Eq. (4) dominant balance yields,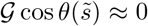 which implies,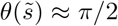, i.e., the snake’s body is approximately vertical, but is incompatible with the boundary condition *θ*(0) = π. In the proximity of,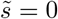, Eq. (4) reduces,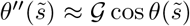 over a length δ determined by the balance 1*/*δ^2^ ∼ 𝒢, i.e., δ ∼ 𝒢 ^*—*1*/*2^. This is in excellent agreement with the numerical results (Fig. 2(c), see SI Sec. S4 for further details of the analysis).

We thus find that the simple choice *m*_a_(*s*) = *B*_a_*κ(s)* increases the effective stiffness of the snake and qualitatively captures the characteristic features of the shape of the snake, i.e. that curvature and muscular activity are large near the lower perch where the snake turns upwards, while in regions where the snake is vertical no muscular forces are required since the gravitational torque vanishes there. However, this choice of *m*_a_(*s*) does not account for the non-local musculature of the snake, and a different choice could potentially accomplish the same task more effectively and might therefore be preferred by the snake. Thus the question arises if there is an *optimal* choice for *m*_a_(*s*), a question we now turn to.

### B. Optimal control of posture

To find the optimal activity profile *m*_a_(*s*) that maintains a standing posture, we need an auxiliary condition, or a cost function, in the spirit of optimal control theory [18, 19]. This function needs to be extremized while being subject to the constraint given by Eq. (2) that determines the shape of the snake with d*m*_a_*/* d*s* being an undetermined control field. Defining the scaled moment,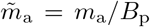, scaled gravity α = ρ*g/B*_p_, and the control,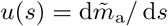, we can define a state vector,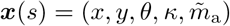 such that its evolution in space can be written as ***x***^′^(*s*) = (cos *θ*, sin *θ*, *n*, α(*l s*) cos *θ u, u*) *f*(***x***(*s*), *u*(*s*)), where we used the definition of,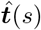 and Eq. (2). The control *u*, and hence the muscular activity, is found by solving the following optimal control problem:

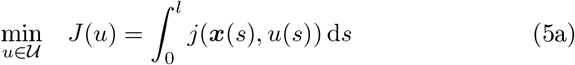

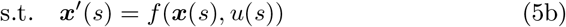

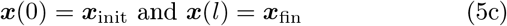

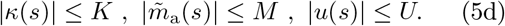

The cost functional *J* in Eq. (5a) is tantamount to the total energy of actuation in terms of the energy density *j*(***x***, *u*), and needs to be minimized over a set 𝒰 of admissible functions. The optimization is constrained by initial and final conditions on ***x*** (Eq. (5c)) as well as bounds on the curvature *κ(s)*, activity,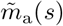 and control *u*(*s*) (Eq. (5d)).

The maximum curvature *κ ∼* 0.5 cm^—1^ is a limit imposed by the snake’s anatomy, and in some experimental trials we observe curvature approaching these literature values for maximal dorsoflexion (see Fig. S4) [20]. We use an order-of-magnitude estimate for *M* from the maximal standing height observed in the experiments, assuming that this is set by limits on muscular force generation, *M ≈* 50 m^—1^ [3, 21] (see SI Sec. S1). Lastly, we set *U* = 100 m^—2^. (See SI Sec. S5 for results with different values.) The boundary conditions are as in Sec. III A, *x*(0) = *y*(0) = 0, *θ*(0) = π, and *θ*(*l*) = *θ*_fin_. We moreover choose *x*(*l*) = —0.1,,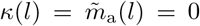, and tune the height of the snake, *y*(*l*) ∈ [0.3, 1.0] to mimic the standing-up process. Finally, we choose the cost or energy density,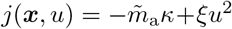, motivated by the idea that the snake strives to minimize the overall energy expanded when assuming a certain posture. The first term is the energy density associated with work done by the active torque,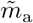 when considering Eq. (2) (with a minus sign to ensure a positive energy) while the second term is added as a regularization term to minimize rapid spatial gradients in the active torque, and is similar in spirit to the quadratic cost used in LQG control, choosing ξ ≪ 1 (specifically, ξ = 0.001). Results for some other choices of *j* are shown in SI Sec. S5, but do not change our results qualitatively.

We numerically solve the optimal control problem using the package `CasADi` [22] and Fig. 3(a) shows the snake posture obtained as the solution. We note the qualitative similarity between our experimental observations and the shapes obtained from our optimal control solutions (see Fig. 3 for three different fractions of the snake being in the air, i.e., *y*(*l*) = 0.4, 0.7, 1.0). Just as for our local kernel, which corresponds to the choice of proportional feed-back in the language of control theory, we find that localizing the active torque in a boundary layer near *s* = 0 is the best strategy for the global optimal control problem. Away from the boundaries the curvature approximately vanishes, i.e, in regions where the snake is almost vertical, active muscular forces are small. However, unlike in the case of local activity treated earlier in Sec. III A, where the curvature at the perch *κ*(*s* = 0) increases as more and more of the snake is in the air, the optimal control problem shows that the perch curvature remains approximately constant during the standing-up process. Interestingly, the muscular activity, 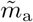(shown in Fig. 3(b)) is approximately equal to the negative curvature,,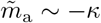 for small values of *l*, while as *l* increases, 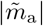 decreases significantly. Furthermore, the optimal control *u*(*s*) takes its limiting value associated with its bound near *s* = 0 for small values of *l* (see Fig. 3(c)) while for larger values of *l* the magnitude of the control decreases significantly. Therefore, muscular actuation and control seem to become less relevant as the standing height increases.

**Figure 3.**
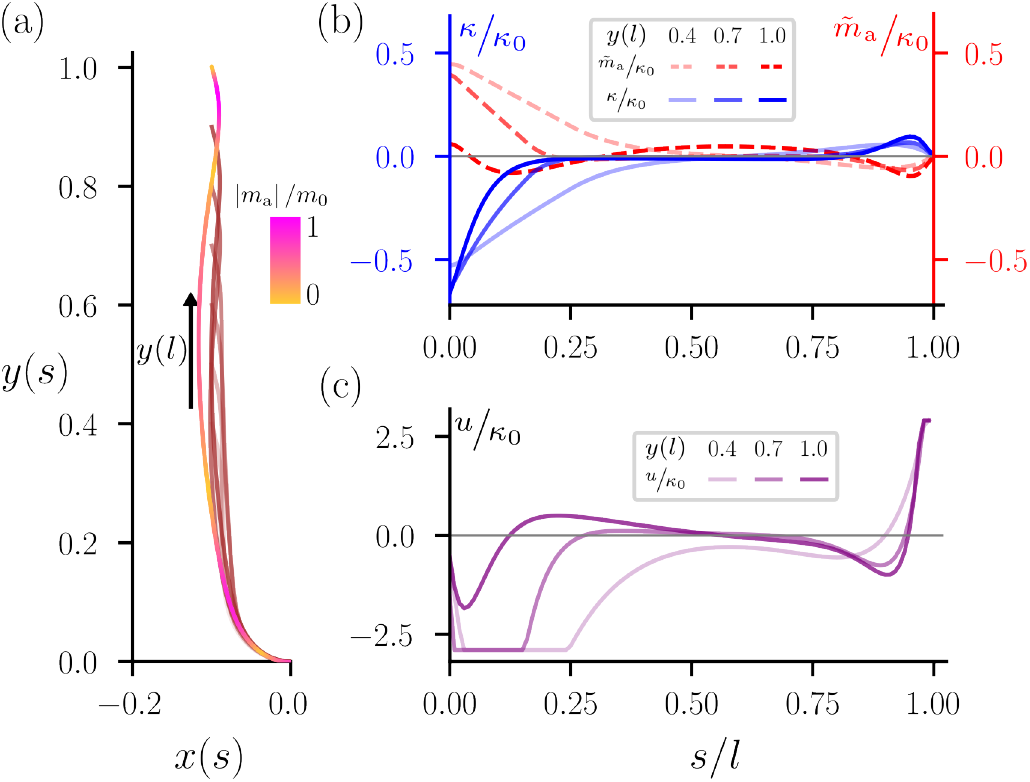
Optimal control of posture. (a) Shapes found for different values of *y*(*s* = *l*) from solving the optimal control problem stated in Eq. (5) and explained in the main text. For the longest curve we show the normalized muscular actuation along the curve using a color map. (b) Curvature *κ(s)* (blue, solid lines) and active moment over bending stiffness,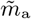 (red, dashed lines) for three values of *y*(*s* = *l*). As before a boundary layer exists near *s* = 0 and away from the boundaries the magnitude of both *κ* and,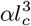 is vanishingly small. We used the same normalization value *κ*_0_ as in Fig. 1. (c) Control *u*(*s*) for the same three values of *y*(*s* = *l*). Note that *u*(*s*) reaches its lower bound for *y*(*s* = *l*) = 0.4, 0.7.

We explain this counter-intuitive behavior as follows: for small *l* the gravitational force is small such that,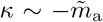 from Eq. (2)^1^. To comply with the boundary conditions, *κ* has to be non-vanishing near *s/l* = 0 and thus,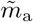 (and *u*) are large. On the other hand, for large values of *l* the gravitational force acting on the snake near the lower perch is much greater. This allows for an actuation profile,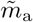 with much smaller overall magnitude that “directs” the gravitational force, such that the resulting energy *J* is minimal while curvature is localized near *s* = 0.

All together, we find that for small *l* the muscular forces are required to bend the shape such that the boundary conditions are fulfilled, while for large *l* the large gravitational forces can be directed towards this goal. By formulating the problem as an optimal control problem we have thus found a much more efficient way of achieving the task, compared with the local approach of prescribing activity proportional to shape as treated earlier in Sec. III A, in a manner reminiscent of classical approaches in control theory, where proportional control is often sub-optimal relative to global optimal control. Our results also suggest that the muscular forces required to achieve a certain posture are not the limiting factor that determines the maximal height a standing snake can reach. Why this might be so is evident from the fact that so far we have considered the standing-up process only as a quasi-static process. However, in our experiment and in the wild, the snake must keep itself balanced as it dynamically pushes its body off the perch. Significant muscular forces are presumably necessary to prevent the snake from falling over, especially at large *l*. Therefore, we must consider the question of *dynamic stability*, which we turn to next.

### C. Postural stability

To better understand the postural stability we consider small perturbations of a given shape. In the absence of any activity, *m*_a_ = 0, the question of stability is equivalent to the classical problem of self-buckling of an elastic rod in a gravitational field [15, 16]. In this limit, the passive elastic filament described by *θ*^′′^(*s*)+α(*l − s*) cos *θ* = 0, where α = ρ*g/B*_p_. For the case of a vertical clamp, i.e., with boundary conditions *θ*(0) = π*/*2, *θ*^′^(*l*) = 0, there is a trivial constant solution *θ* = *θ*_0_ = π*/*2. This solution, however, can be unstable when the length *l* is large enough. To see this, we introduce a small perturbation ϕ, *θ* = *θ*_0_ + ϕ, with ϕ ≪ 1, leading to a linearized equation for ϕ (*s*) that has non-trivial solutions if,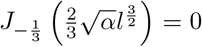 where *J*_*n*_ is the Bessel function of the first kind [16]. This classical (Greenhill) criteria for self-buckling yields,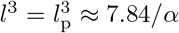 such that for *l > l*_p_ the solution *θ* = *θ*_0_ is unstable.

We now investigate the effect of the presence of an active moment on the stability. If we choose a local proprioceptive feedback for the active moment with *m*_a_(*s*) = *B*_a_*κ(s)*, the active moment effectively simply renormalizes the bending elasticity leading to the same stability criterion as in the passive case, but with *B*_p_ ⟶ *B*_p_ + *B*_a_. Therefore, for *B*_a_ > 0 the effective stiffness of the snake becomes larger and the critical length increases as well, since,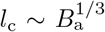. However, in reality such an ideal local feedback is difficult to achieve due to the complex musculature of the snake that spans multiple vertebrae [13]. To investigate how the presence of non-local feedback impacts the stability, and if it can perhaps even be utilized to increase the critical length, we assume a non-local feedback of the form:

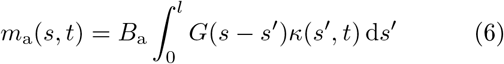

with a Gaussian kernel,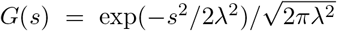 and a new characteristic length scale λ. We note that in the limit λ ⟶ 0 this kernel converges towards the delta distribution and we recover the case of local feed-back where *m*_a_(*s*) = *B*_a_*κ(s)*. To investigate the effect of non-locality on the stability we numerically compute the bifurcation diagram for Eq. (2) with the choice given by Eq. (6) for several values of λ and various *B*_a_*/B*_p_ in order to compare the stability with both the local feed-back and the passive cases (see Materials and Methods for details).

In Fig. 4(a), we show the maximum deviation from the initial vertical configuration *θ*_max_ = max | *θ*(*s*) − π*/*2 | for *B*_a_*/B*_p_ = 0.5. We find that the presence of non-locality results in an earlier onset of instability (at smaller values of α*l*^3^) compared with the local feedback case (λ = 0), implying that the non-locality hampers/reduces the stability. When λ ≳ *l*_p_, i.e., the snake musculature spans a length comparable to the Greenhill length, the active moment becomes negligible and we approximately recover the passive case (*m*_a_ = 0) since the filament is self-stabilized by long-range musculature; indeed in the limit λ ⟶ ∞ the Gaussian feedback kernel in Eq. (6) vanishes and *m*_a_ ⟶ 0. The critical length,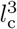 for self-buckling decreases monotonically with increasing λ (see Fig. 4(b)), a trend consistent for different values of *B*_a_*/B*_p_, and all the bifurcation curves converge towards the passive curve *B*_a_ = 0 for λ *≳ l*_p_ (see Fig. 4(b)).

**Figure 4.**
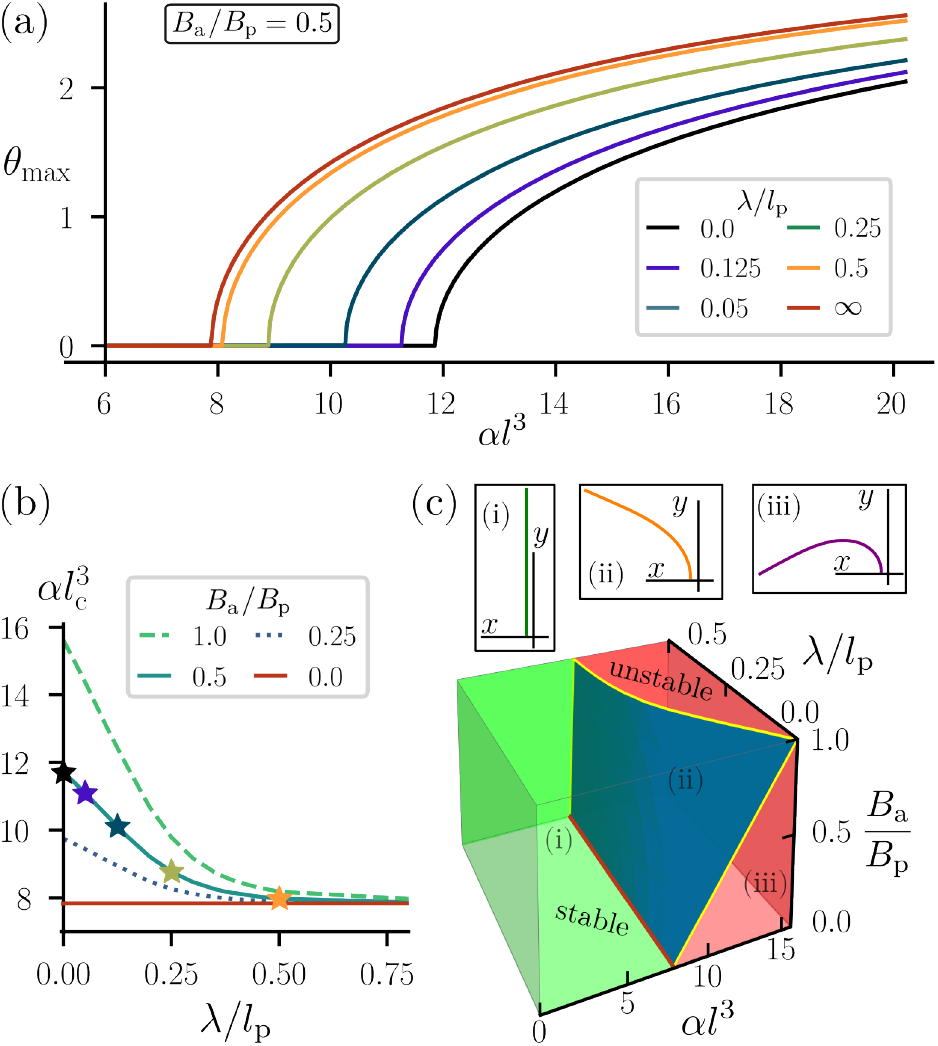
Stability of standing. (a) Bifurcation diagram for non-local feedback with Gaussian kernel (Eq. (6)) with standard deviation *λ. *θ**_max_ = max |*θ*(*s*) *— π/*2| corresponds to the largest deviation of the beam from its initial straight configuration. The left-most (*λ* = ∞) and right-most (*λ* = 0) curves correspond to the passive *B*_a_ = 0 and the local case *m*_a_ = *B*_a_*κ*, respectively. *l*_p_ is the critical length in the passive case. (b) The critical parameter,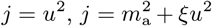 as a function of non-locality *λ* for different values of *B*_a_*/B*_p_. The stars indicate the critical values found in (a) for the *λ*-values considered there. The solid orange line highlights the case of vanishing muscular activation, *B*_a_, where the equations reduce to the classical self-buckling problem. All curves converge to this values as *λ* → ∞. (c) Stability phase diagram for different values of *λ* and *B*_a_. Points in space to the right of the blue surface correspond to unstable configurations, while points to the left correspond to stable ones. Three example configurations from the stable and unstable region are shown in the inset. (i) is a stable pose, (ii) and (iii) are unstable poses, corresponding to (*al*^3^, *B*_a_*/B*_p_, *λ/l*_p_) = (2, 0.8, 0.1), (12, 0.5, 0.25) and (15, 0.1, 0.1), respectively.

A three-dimensional stability phase diagram summarizes our results and is shown in Fig. 4(c), where we illustrate the stable and unstable phases as a function of all

three free parameters in the model, namely the (scaled) non-locality parameter λ*/l*_p_, the relative strength of the active moment *B*_a_*/B*_p_, and the (scaled) weight of the snake α*l*^3^. We find that: (*i*) the boundary between the stable and unstable regions is approximately linear for all values of λ (see Fig. S9 in SI for further details), but with a varying slope and offset such that for large values of λ the boundary converges towards the vertical α*l*^3^ ≈ 7.84; (*ii*) the stable region is largest for λ = 0 but stability persists in the presence of non-linearity. A similar phase diagram but for a biologically relevant parameter range is presented in SI Fig. S9 and allows for similar conclusions.

## IV. CONCLUSIONS AND OUTLOOK

How do our different theoretical frameworks connect back to our experimental observations? Since we were unable to directly measure the muscular activity during our experiments we can only indirectly estimate *m*_a_(*s*). Assuming local feedback for simplicity, we can use the expression for *l*_c_ to obtain an order-of-magnitude estimate of the maximal muscular activation from the largest vertical distance *L* that the snakes were able to bridge in the experiments. We find,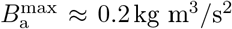 (see SI Sec. S1) such that 𝒢 ^max^ ≈7.8. This is significantly lower than the parameter values used in the numerical solutions in Sec. III A, especially since,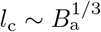. Furthermore, we recall that the magnitude of the optimal active moment found in Sec. III B *decreases* as *l* increases. This results in an overall mismatch that demands explanation.

To this end, we recall that in Secs. III A, III B we considered only the static posture with no regard for its stability, while above we estimated,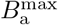 from stability considerations. This leads us to the conjecture that the muscular forces required to stabilize a given posture are much larger than the forces required to establish it (at least once *l/l*_tot_ is sufficiently large). Said simple, it is stability that limits the maximal standing height. While more experiments and quantitative measurements of the muscular activity are required to verify this hypothesis, there is qualitative evidence of this in our observations. Indeed, for small lengths of the snake in the air the posture is approximately static, while for larger lengths the snakes sway slowly in a manner suggestive of the dynamic stabilization of an inverted (soft) pendulum, see SI Movies 1-4. The amplitude of these oscillations are relatively small, but clearly present. We speculate that these are a result of the stabilization process requiring large muscular forces.

Our study was inspired by an extreme in postural control exhibited by a slender organism, arboreal snakes such as the brown tree snake and the scrub python that lift themselves into an almost upright position with up to 70% of their body length into the air. To quantify the muscular control of the posture, we introduced a minimal model of an active filament and considered a variety of models for activity and showed that they were sufficient to qualitatively reproduce the key experimental observations associated with the formation of a boundary layer near the lower perch, where curvature is large [13], and a vanishingly small curvature away from this region. The two possibilities mentioned in the introduction, namely supporting the body by activating antagonistic muscles to increase overall stiffness and activating only some muscles to achieve a particular shape, correspond to our results in Sec. III A and Sec. III B, respectively. In both cases, the required effort to achieve a given posture can be reduced by localizing muscular activity near the perch, while maximizing the total vertical length of the body in the air, allowing for efficient navigation in arboreal environments with vertical gaps comparable with the total body length. Though both strategies, a local proportional activity and a globally optimal one, can be used to achieve the observed shape, the latter is more energy efficient and thus is likely to be chosen by the snake. However, this prediction needs to be tested by future experiments. Further, we found that the presence of muscular activity increases stability against a self-buckling instability relative to the passive case, and that this increase in stability persists even if the muscular feedback is non-local.

Our framework is likely to be applicable to a range of other similar situations, e.g., the magnificent posture of a cobra, eels in flows [4], elongated appendages of other animals [17, 23], plant shoots [24–26], and a range of animal and plant-inspired soft robotic mimics [27–30]. Indeed, the phase diagram Fig. 4(c) predicts a general relation between stability and (non-)local active forces which can be used to guide robotic designs such as the development of slender medical devices (like endoscopes) [31–34]. Lastly, our approach to integrate optimal control theory to study posture and its stability or the effect of non-locality in space (and time) on the feedback response is likely to carry over more broadly to neuromechanical systems.

## ACKNOWLEDGEMENTS

This work was supported by the NWO Rubicon grant (L.A.H.), the Simons Foundation (L.M.) and the Henri Seydoux Fund (L.M.).

## MATERIALS AND METHODS

The code necessary to reproduce all the results will be made available upon publication.

The numerical solutions shown in Sec. III A were found using `Auto-07p` [35]. To solve the optimal control problem we use the non-linear programming solver of `CasADi` [22], using the Ipopt software package with 100 discretization steps and an interpolating polynomial of degree 3. We solve the optimization problem for twenty values of *l* [1.01 *y*(*l*), 1.2 *y*(*l*)] and among the cases where a solution is found we select the one with minimal ∫ |*m*_a_ | d*s*.

To investigate the stability in the case of non-local feed-back we solve the equation of motion

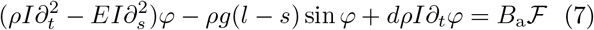

with boundary conditions ϕ (0) = *a*ϕ^′^(0), ϕ^′^(*l*) = 0, *a* = 10^—6^, and *d* = 0.5. For local feedback,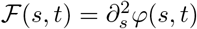, for non-local feedback ℱ (*s, t*) = (*G*^′^ ⋆ ϕ^′^) (*s, t*), with *G* the Gaussian kernel defined in the main text. Integrating by part, it is possible to write the latter as

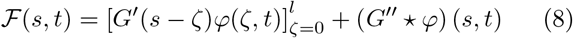

which is the expression we use to numerically solve the equations for efficiency reasons. The convolution is computed using a Simpson’s rule FFT method described in Ref. [36]. We solve the equations using methods of line using the ODE solver of `Julia` with 2^8^ + 1 spatial discretization points.

## AUTHOR CONTRIBUTIONS

L.M. conceived of the study. L.A.H., P.B., I.C.D., S.G.P., and L.M. carried out theoretical and computational work, B.C.J. performed experiments. All authors wrote, reviewed, and edited the manuscript.

## Supplementary Information

### S1. ESTIMATE OF EXPERIMENTAL PARAMETERS AND LIST OF PARAMETERS

To estimate the parameters for the snake we proceed as follows. We stress that the following is only an order-of-magnitude estimate of the parameters. The density of the brown tree snake is calculated by dividing a total mass of a snake by its total length, thus assuming that the density is constant along the snake’s body. Averaging the density of six different snakes we find ⟨ ρ⟩ = 0.36 ± 0.03 kg/m.

We observe that just after touching the target perch the snake appears to relax and starts oscillating (see Movie 5). We use the frequency of this oscillation to roughly estimate the passive bending stiffness by modeling the process as an oscillating beam. The fundamental frequency of a beam of length *l* and stiffness *B*_p_ is

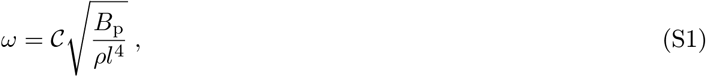

where the constant 𝒞 is set by the choice of boundary conditions. For a simply supported beam 𝒞 = π*/*2. From movie Movie 5 we find ω ≈ 0.8 Hz and *l* = 0.7 m such that

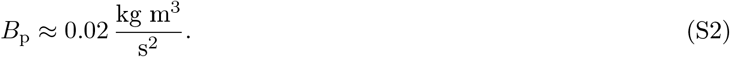

As described in the main text, the critical length of the buckling instability is given by

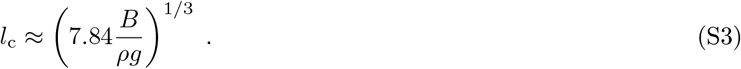

Were there not muscular actuation, *B* = *B*_p_ and *l*_p_ ≈ 0.36 m. Assuming that the largest observed standing height corresponds to the largest possible muscular activation, we can estimate an upper bound for the active bending modulus from setting *B* = *B*_p_ + *B*_a_ where we assumed local feedback *m*_a_ = *B*_a_*κ* such that the effective stiffness is given by the sum of passive and active component. With *l*_c_ ≈ 0.8 m we have

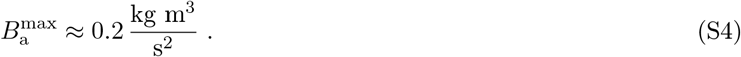

We can compare our estimate with literature results from *in vivo* measurements of stress generation of limb muscles of lizards. It was found that the maximal generated stress is about σ ≈ 200 kN m^—2^ [1, 2] such that *B*_a_ ∼ π^2^σ*R*^4^ ≈ 0.1 kg m^3^*/*s^2^, approximating the snake as a cylinder of constant radius *R* = 3 cm. These two independent ways of estimating the maximal muscular actuation thus agree in order-of-magnitude.

Finally, in Tab. S1 we list the variables used in the main text together with their respective units.

### S2. IMAGE ANALYSIS

To measure the shape and curvature of the snakes in the experiments during the standing-up process we proceed as follows. A video starts being recorded after the snake is put on the lower perch. From the recording we extract all frames for the time between when the snake’s head first leaves the lower perch until the point where the snake reaches the upper perch. An example of a frame is shown in Fig. S1(a). For each frame we separate the snake from the background using a personalized `Python` script using the skimage library. An example of the resulting image is shown in Fig. S1(b). Having isolated the shape of the snake, we identify the epaxial edge of this shape. The resulting curve is interpolated such that we obtain a smooth curve that approximates the shape of the snake well (see Fig. S1(c)). This interpolated curve in turn is used to compute the local curvature at each point, obtaining a function *κ(s)*. Performing this procedure for each frame, we obtain a time evolution of the curvature. For the results presented in this paper an experiment for a given snake and height *L* is repeated several times such that we can obtain an averaged curvature evolution. More details on this procedure can be found in Sec. S3 below. For the three brown tree snakes (snout-vent lengths [SVL] 172 cm, 169 cm, and 120 cm) we analyzed in total one trial for which the horizontal gap was 40% of the SVL, ten trials at 45% and seven trials at 50%. For the python (SVL 102 cm) we analyzed four trials at 29% SVL, one trial at 49%, 2 trials at 66% and four trials at 70%.

### S3. SHAPE EVOLUTION OF STANDING SNAKES

The figure analogous to Fig. 1 (where we showed snapshots and curvature for the scrub python) in the main text but now for the brown tree snake is Fig. S2. As for described in the main text, we record a video of the snake from a lateral and a dorsal point of view (snapshots in Fig. S2(a, b)). An overlay of different snapshots covering the entire process is shown in Fig. S2(c). The resulting lateral curvature at different points during the process is shown in Fig. S2(d). As mentioned in the main text, these results are averages over several experimental runs (four in this case) with the same experimental setup and the same snake. We will explain in more detail the averaging process below.

**Table S1.**
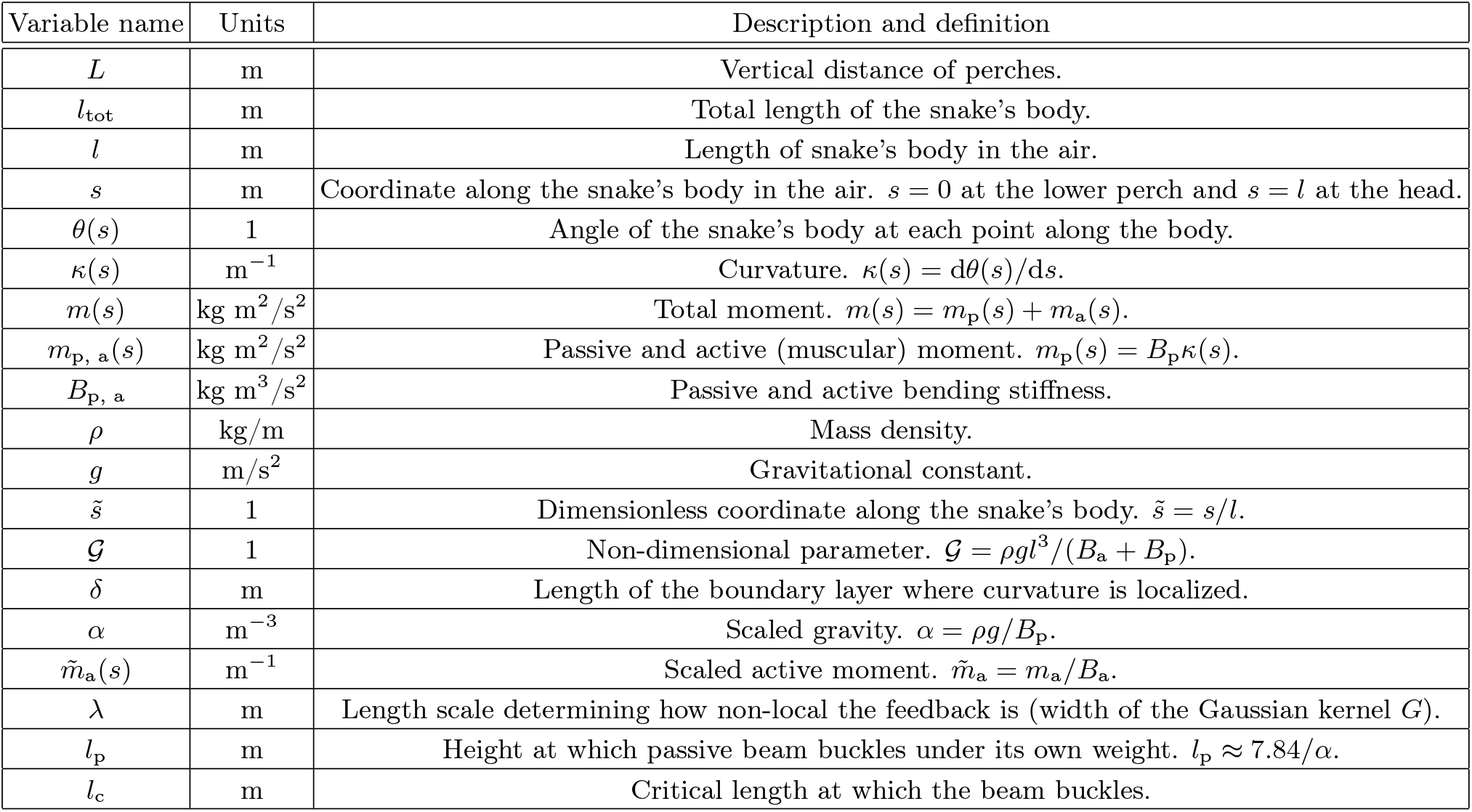
List of Variables.

| Variable name | Units | Description and definition |
| --- | --- | --- |
| $L$ | m | Vertical distance of perches. |
| $l_{\text{tot}}$ | m | Total length of the snake's body. |
| $l$ | m | Length of snake's body in the air. |
| $s$ | m | Coordinate along the snake's body in the air. $s = 0$ at the lower perch and $s = l$ at the head. |
| $\theta(s)$ | 1 | Angle of the snake's body at each point along the body. |
| $\kappa(s)$ | $\text{m}^{-1}$ | Curvature. $\kappa(s) = d\theta(s)/ds$. |
| $m(s)$ | $\text{kg m}^2/\text{s}^2$ | Total moment. $m(s) = m_p(s) + m_a(s)$. |
| $m_{p, a}(s)$ | $\text{kg m}^2/\text{s}^2$ | Passive and active (muscular) moment. $m_p(s) = B_p \kappa(s)$. |
| $B_{p, a}$ | $\text{kg m}^3/\text{s}^2$ | Passive and active bending stiffness. |
| $\rho$ | $\text{kg/m}$ | Mass density. |
| $g$ | $\text{m/s}^2$ | Gravitational constant. |
| $\tilde{s}$ | 1 | Dimensionless coordinate along the snake's body. $\tilde{s} = s/l$. |
| $\mathcal{G}$ | 1 | Non-dimensional parameter. $\mathcal{G} = \rho g l^3 / (B_a + B_p)$. |
| $\delta$ | m | Length of the boundary layer where curvature is localized. |
| $\alpha$ | $\text{m}^{-3}$ | Scaled gravity. $\alpha = \rho g / B_p$. |
| $\tilde{m}_a(s)$ | $\text{m}^{-1}$ | Scaled active moment. $\tilde{m}_a = m_a / B_a$. |
| $\lambda$ | m | Length scale determining how non-local the feedback is (width of the Gaussian kernel $G$). |
| $l_p$ | m | Height at which passive beam buckles under its own weight. $l_p \approx 7.84/\alpha$. |
| $l_c$ | m | Critical length at which the beam buckles. |

**Figure S1.**
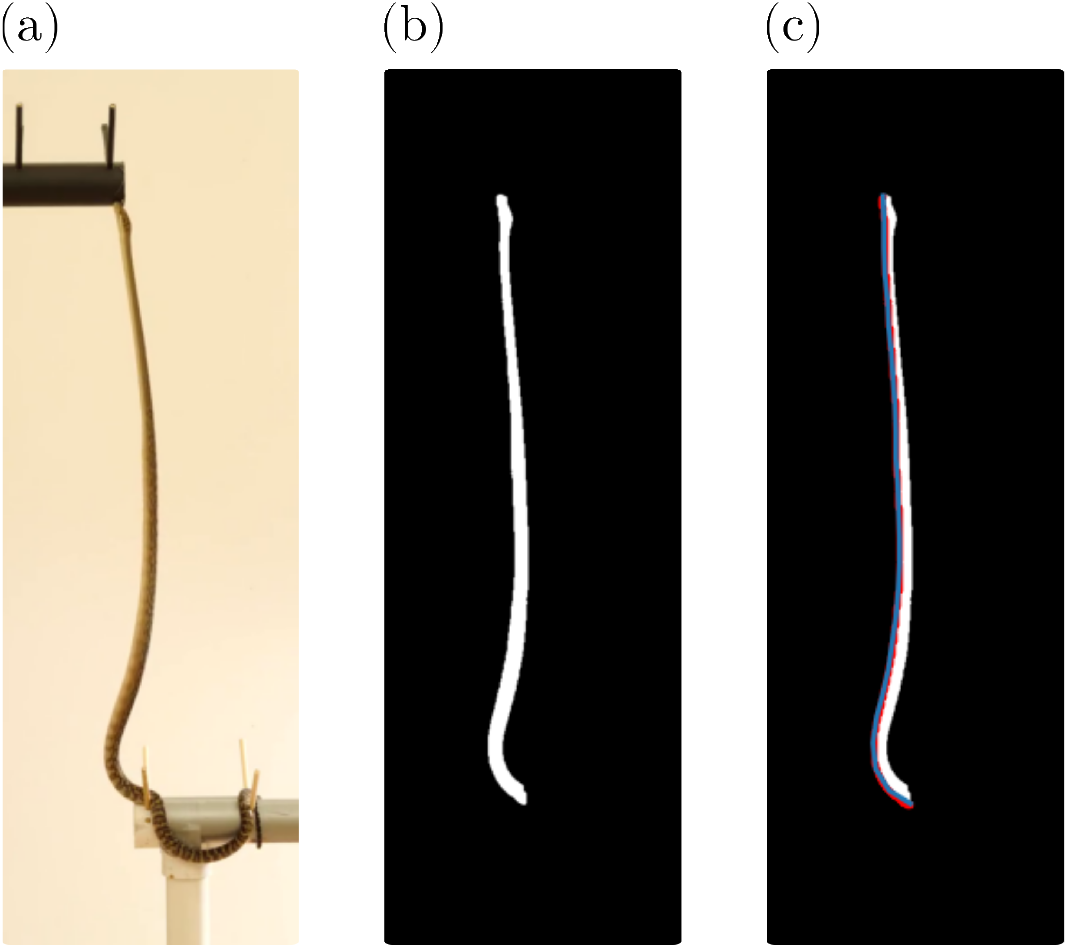
Curvature measurement. (a) Example of a single frame from a video recording of the standing-up process. (b) Removal of background and isolation of the shape of the part of the snake that is not in contact with the perch. (c) We first identify the epaxial edge of the shape (here shown as a red line) and then find a smooth interpolating function to approximate this curve.

Throughout this article we focus on the shape observed from the lateral view, such as Fig. S2(a) and measure curvature in this plane. The shape observed from the dorsal view is not trivial either, and in particular the curvature in this plane does not vanish, as can be easily seen in Fig. S2(b) for example. However, we find that the shape in this plane is not universal, and that the curvature occurs both with negative and positive sign. On the other hand, the later shape is seen to be qualitatively similar across all experiments. This is illustrated for a few examples in Fig. S3. Thus, when comparing many runs, the lateral curvature is conserved while the dorsal curvature is not. For simplicity, we therefore focus on the more universal lateral dynamics. The dorsal shape seems to be intimately connected with the specific manner in which the snake is position and moving on the perch before standing up. However, we have not investigated this relation in detail.

**Figure S2.**
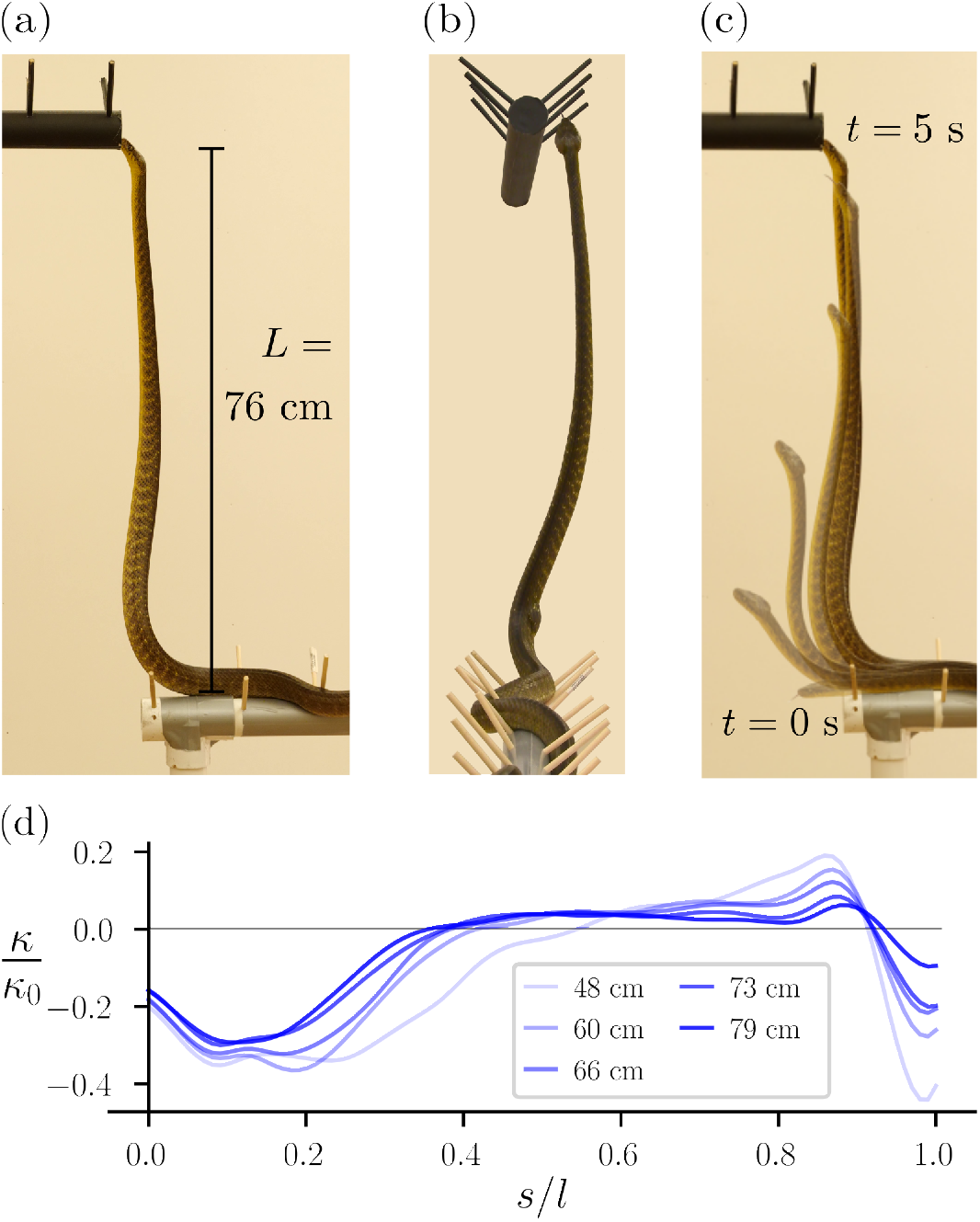
Posture of standing brown tree snake. (a) Lateral and (b) dorsal view of a brown tree snake which is standing on a lower perch, with the upper perch at a vertical distance *L* = 0.76 m. Shown is the position shortly before the snake makes contact with the upper perch. (c) Overlay of the brown tree snake at different time steps, with earlier times being more transparent. The first shown position is the frame where the snake’s head first leaves the perch while the last shown position is the last frame before the snake makes contact with the upper perch. Time between subsequent shown positions is about 0.75 sec. (d) Measured curvature *n* along a brown tree snake’s body at different times during the climbing process. The length of the snake’s body that is in the air increases from *l* = 0.48 m to *l* = 0.79 m. Curvature is normalized by *κ*_0_ = 1*/*3.4 cm, the inverse width of the snake. The gap size *L* = 0.86 m at which these experiments were performed corresponds to about 50% of the snake’s total length.

The curvature extracted from the experiments and presented in Fig. 1 (scrub python) and Fig. S1 (brown tree snake), respectively, are averages over several experimental trials. In either case, the same experimental setup (same height *L*) was used and the same snake was recorded several times standing up. In both Fig. 1 and Fig. S1 the curvature shown is an average over four trials each. In Fig. S4 we show the average of three curves together with the standard deviation. We find that in both cases the standard deviation at late times during the process is significantly smaller than at earlier times. As mentioned in the main text, even when using the same experimental setup and the same snake, the total time between the snake first lifting off the lower perch and the snake first touching the upper perch varies between different trials. Thus, to compare different trials and to take the average, we use the length *l* of the snake’s body that is in the air as a measure of progress and a proxy of time. That is, for each video of a given *L* and snake we choose a few values for *l* and pick the closest corresponding frame for each of the (in this case four) videos. Note that due to the limited frame rate (30 frames/sec) it is not always possible to pick a frame that exactly corresponds to the length *l*. Thus we pick the frame that is closest to this value. This results in an error in the length *l* associated with the different curves in Figs. 1, S1, S4. However, we find that this error is small (the largest relative length difference we found was ≈ 0.02) and thus it is not relevant. For completeness we report the errors in Tab. S2.

**Table S2.** Error of *l*. The error associated with *l* when averaging over four experiments for the scrub python (Fig. 1) and the brown tree snake (Fig. S2), respectively.

|  |  |  |  |  |  |
| --- | --- | --- | --- | --- | --- |
| Scrub python | $29 \pm 0.2$ cm | $39 \pm 0.2$ cm | $49 \pm 0.1$ cm | $59 \pm 0.3$ cm | $69 \pm 0.4$ cm |
| Brown tree snake | $48 \pm 1.1$ cm | $60 \pm 0.4$ cm | $66 \pm 0.7$ cm | $73 \pm 0.6$ cm | $79 \pm 0.5$ cm |

**Figure S3.**
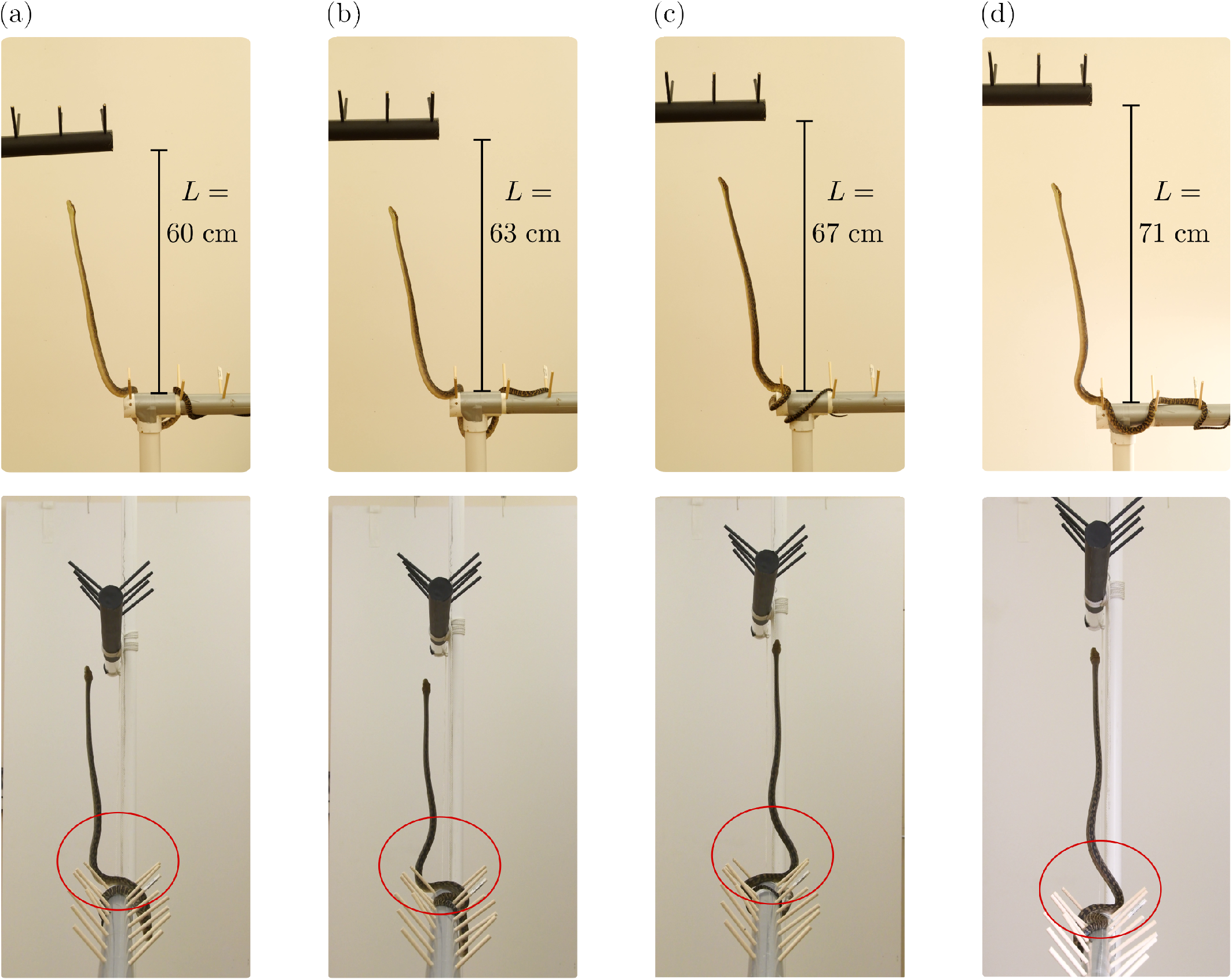
Lateral and dorsal curvature. Several snapshots of the standing scrub python for different values of vertical distance *L*, namely (a) *L* = 0.60 m, (b) *L* = 0.63 m, (c) *L* = 0.67 m, (d) *L* = 0.71 m. The top row of snapshots show the lateral view while the bottom row show the dorsal view. In the bottom row we highlighted with a red circle the region of largest curvature. While the lateral shape is very similar for all four snapshots, the dorsal shape is not.

**Figure S4.**
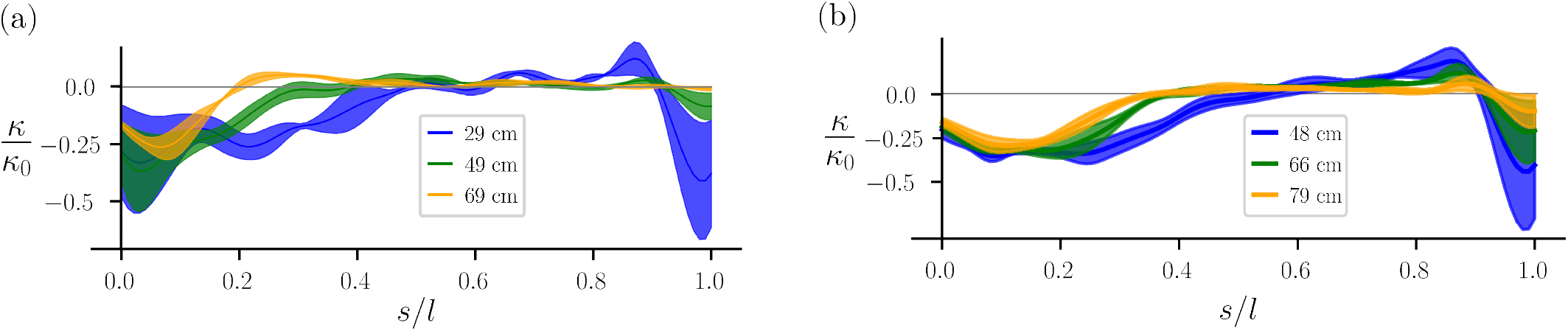
Averages and standard deviations of curvature. We show in (a) the averages (solid line) as well as the associated the standard deviation (shaded area) of the curvature *κ(s)* for three values of *l* for the scrub python. The average curves are the same as the ones shown in Fig. 1 for the corresponding values of *l*. In (b) we present the same data for the brown tree snake, cf. Fig. S2.

**Figure S5.**
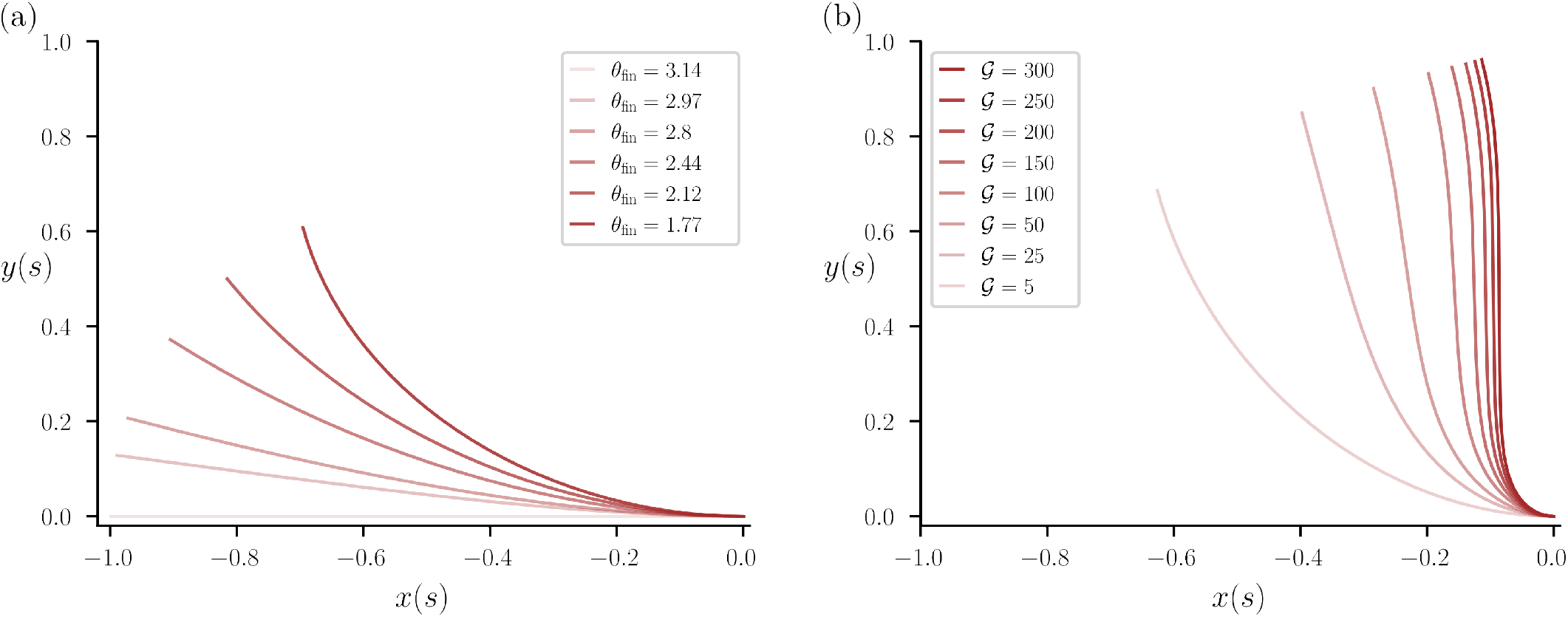
Lift-up and stand-up with proportional feedback. (a) We model the lift-up behavior by numerically solving Eq. (4), keeping 𝒢 = 1 constant here but for different values of *θ* (*s* = *l*) = *θ* _fin_. (b) Results for the stand-up phase, that is the same data as in Fig. 2(a) in the main text but with all curves now normalized to be of length *l* = 1.

### S4. POSTURE WITH LOCAL FEEDBACK

While the lift-up phase is seen to not be as universal as the stand-up phase, we nevertheless can model a simple, idealized version of this dynamics. Namely, we change the angle *θ* (*s* = *l*) = *θ*_fin_ near the head while keeping constant, i.e., assuming that the length of the body that is in the air is approximately constant during the lift-up phase. The resulting solutions for some values of *θ*_fin_ are shown in Fig. S5(a). The final value of *θ*_fin_ is the one that is kept constant in the standing-up phase in the main text. To illustrate the change of geometry during this phase more clearly, we present in Fig. S5(b) the same data as in Fig. 2(a) in the main text, but with all curves now having the same length *l* = 1. This highlights especially the change in shape for small values of 𝒢.

As we have mentioned in the main text, there exists a boundary layer in the shape of snake close to the branch at,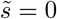 and the equilibrium equation governing the shape of the snake is,

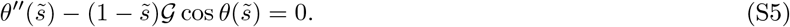

This reduces to,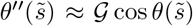 near the branch i.e.,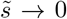, which needs to be solved with boundary conditions *θ* (0) = π to *θ* (1) = π*/*2 in order to capture its shape. We can see that upon scaling,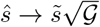, we get,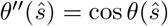.This system is analogous to the shape of hair near the scalp as discussed in Ref. [3] and has the solution,

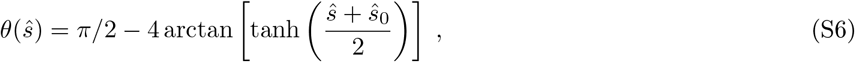

with,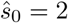 artanh[tan(π*/*8)]. From this solution we can find the curvature,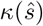 to be

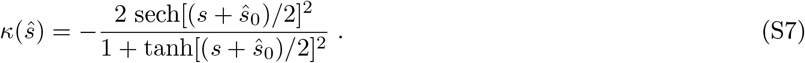

Note that since,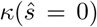 is constant, we immediately find from rescaling,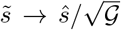 that,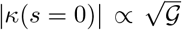. In Fig. S6 we show a comparison of the analytical result Eq. (S7) with simulation results.

**Figure S6.**
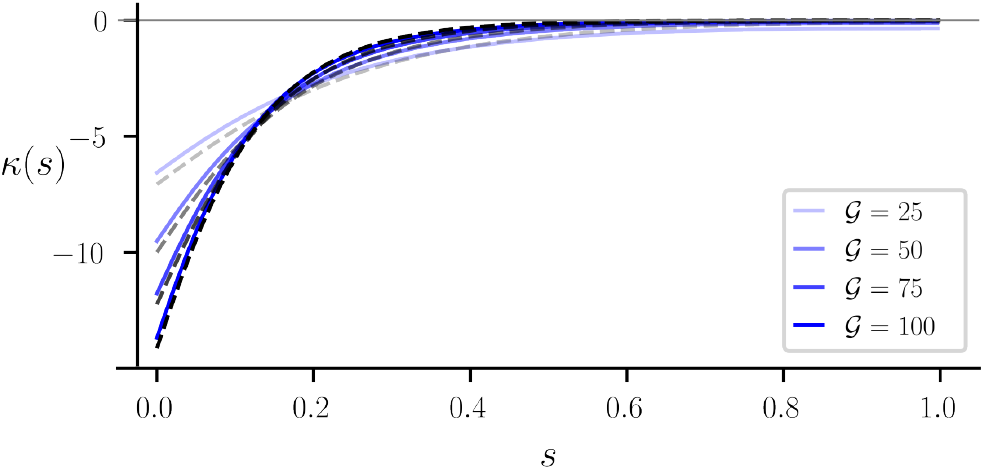
Comparison of results from simulation and analytics. The blue, solid lines shown here are the curvature curves found from the simulations assuming local feedback for different values of 𝒢, i.e., similar to Fig 2 in the main text. The black, dashed lines are the analytical results for the same values of 𝒢 from Eq. (S7).

### S5. OPTIMAL CONTROL THEORY

In this section we present results for the posture found using optimal control theory similar to the ones shown in the main text, but first for different parameter values and second for different cost functions. In Fig. S7 we show the shape, muscular actuation, curvature, and control that we find for solving the optimal control problem for different values of α = ρ*g/B*_p_ and choosing different bounds for the control *u*. We furthermore stress again that, as in the main text, the length *l* is undetermined and the optimal control problem has a solution not just for one value of *l*. Therefore, we solve the problem for various different values of *l* (in this case 20 values in the region [1.01 *y*(*l*), 1.3 *y*(*l*)]) and pick as the presented solution the case for which the integrated muscular activation, ∫ | *m*_a_ | d*s*, is minimal. In most cases this corresponds to picking the smallest value of *l* for which the problem is solvable. Note that in Fig. S7(a) there are no gravitational forces, α= 0, such that,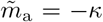 exactly. For larger values of α the curves diverge from this relationship, in particular near *s* = 0 and for large *l* when gravitational effects are the strongest.

**Figure S7.**
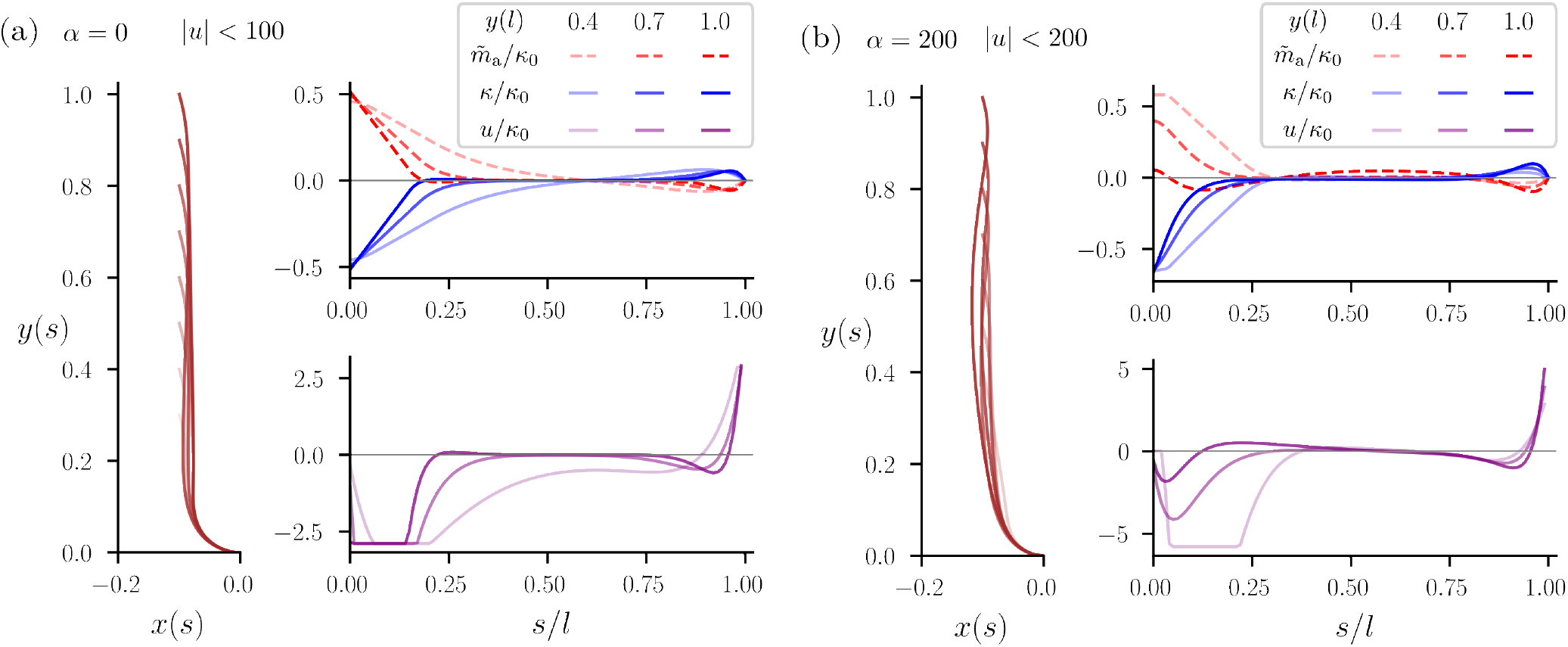
Posture from optimal control theory for different parameters. We solve the optimal control problem Eq. (5) in the main text for different values of α and |*u*|. All other parameters are the same as in the main text. In panel (a) α= 0, |*u*| *<* 100 and in (b) α = 200, |*u*| *<* 200.

To investigate the effect of the cost function *j* on the shape and control, we solved the optimal control problem for one set of parameters for three different choices of *j*, namely,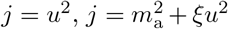, and *j* = − *m*_a_*κ* + ξ *u*^2^ (the same as above and in the main text). As described in the main text, we add ξ *u*^2^ as a regularization term for the second and third cost function, with ξ a small value, here ξ = 0.001. The results are shown in Fig. S8 for two choices of *y*(*s* = *l*).

**Figure S8.**
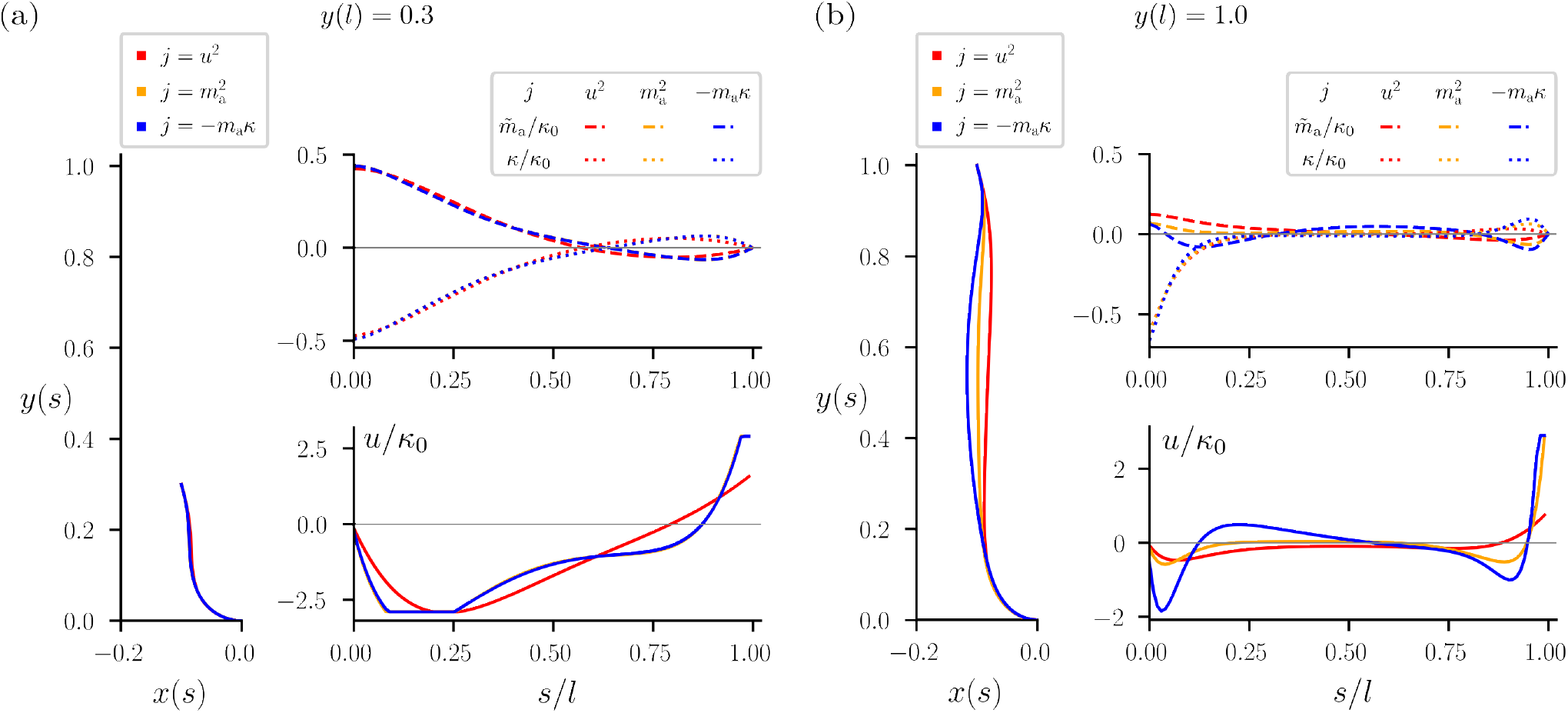
Posture from optimal control theory for different cost functions. We solve the optimal control problem Eq. (5) for the same parameter values as in the main text. However, we here solve the control problem for three different values of the cost function *j*, namely,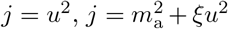, and *j* = *—m*_a_*κ* + ξ*u*^2^ (the same as above and in the main text). In (a) we show the results for *y*(*l*) = 0.3 while in (b) we set *y*(*l*) = 1.0. Note that the blue,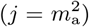 and yellow (*j* = *—m*_a_*κ*) curves are almost identical for small *y*(*l*) (where *m*_a_ *∼ k*) while for larger *y*(*l*) they differ.

### S6. NON-LOCAL FEEDBACK

For the non-local feedback investigated in the main text we can create two-dimensional bifurcation phase diagrams that are essentially λ = const. slices of the three-dimensional diagram presented in Fig. 4(c). We show two examples in Fig. S9(a, b) to present the geometry of the phase-separating boundary more clearly.

**Figure S9.**
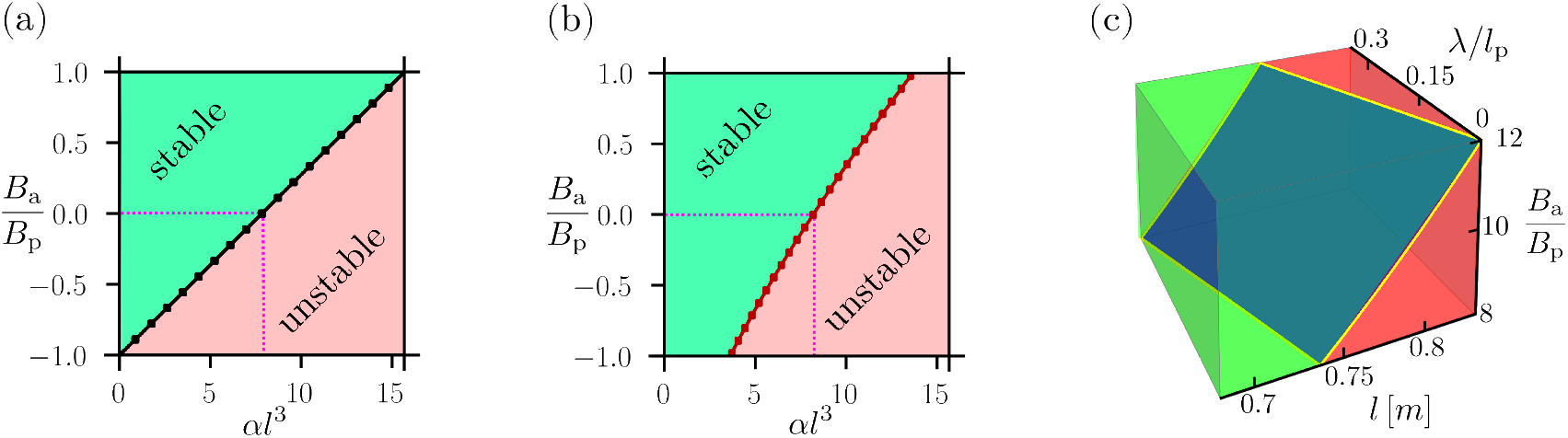
Stability phase diagram for non-local feedback. We numerically solve the stability problem for the non-local Gaussian kernel with standard deviation λ as described in the main text. In (a) we set λ = 0 and thus recover the local feedback with the pink dashed line highlighting the passive case. In (b) λ = 0.5 and we find that the boundary between stable and unstable region is still approximately linear, but with a different slope and offset. In (c) we present three-dimensional stability phase diagram for values of the parameters estimated from the experiments (see Sec. S1). A small but finite value of λ is seen to not significantly shift the boundary of stability.

Lastly, we consider the stability for values close to the values estimated from the experimental observations, see Sec. S1. Namely, with *l*_p_ ≈ 0.36 m and,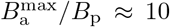 we find the stability diagram shown in Fig. S9(c). We estimated,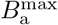 assuming purely local feedback. In Fig. S9(c) we shown how the stability is affected by a small value of non-locality, λ *<* 0.3*l*_p_. This non-locality could be due to, e.g., the finite size of the snake’s muscles.

### S7. MOVIE DESCRIPTIONS

**Movie 1**. The movie shows a scrub python from both the lateral and dorsal point-of-view while it is standing up, with the gap size being *L* = 50 cm.

**Movie 2**. The movie shows a brown tree snake from both the lateral and dorsal point-of-view while it is standing up, with the gap size being *L* = 50 cm.

**Movie 3**. The movie shows a scrub python from both the lateral and dorsal point-of-view while it is standing up, with the gap size being *L* = 71 cm. Additionally, the lateral curvature *κ(s)/n*_0_ along the snake at each time frame is shown.

**Movie 4**. The movie shows a brown tree snake from both the lateral and dorsal point-of-view while it is standing up, with the gap size being *L* = 86 cm. Additionally, the lateral curvature *κ(s)/n*_0_ along the snake at each time frame is shown.

**Movie 5**. The movie shows a scrub python from dorsal point-of-view while it is standing up, with the gap size being *L* = 77.5 cm. Towards the end of the process, large oscillations can be observed. Furthermore, the snake visibly relaxes once it comes in contact with the upper perch.

## Footnotes

1 In fact, in the limit α = 0 we find,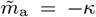 exactly, see SI Sec. S5.

